# A Haptoglobin (HP) Structural Variant Alters the Effect of *APOE* Alleles on Alzheimer’s Disease

**DOI:** 10.1101/2022.09.22.508749

**Authors:** Haimeng Bai, Adam C. Naj, Penelope Benchek, Logan Dumitrescu, Timothy Hohman, Kara Hamilton-Nelson, Asha R. Kallianpur, Anthony J. Griswold, Badri Vardarajan, Eden R. Martin, Gary W. Beecham, Jennifer E. Below, Gerard Schellenberg, Richard Mayeux, Lindsay Farrer, Margaret A. Pericak-Vance, Jonathan L. Haines, William S. Bush, Alzheimer’s Disease Genetics Consortium

## Abstract

**Background:** Haptoglobin (HP) is an antioxidant of apolipoprotein E (APOE), and previous reports have shown HP binds with APOE and amyloid-β (Aβ) to aid its clearance. A common structural variant of the *HP* gene distinguishes it into two alleles: *HP1* and *HP2*.

**Methods:** *HP* genotypes were imputed in 29 cohorts from the Alzheimer’s Disease (AD) Genetics Consortium (N=22,651). Associations between the *HP* polymorphism and AD risk and age of onset through *APOE* interactions were investigated using regression models.

**Results:** The *HP* polymorphism significantly impacts AD risk and age at onset in European-descent individuals (and in meta-analysis with African Americans) by modifying both the protective effect of *APOEε2* and the detrimental effect of *APOEε4*, especially for *APOEε4* carriers.

**Discussion:** The effect modification of *APOE* by *HP* suggests adjustment and/or stratification by *HP* genotype is warranted when *APOE* risk is considered. Our findings also provided directions for further investigations on potential mechanisms behind this association.

## 1 INTRODUCTION

Alleles of the *APOE* gene are the strongest genetic factor for sporadic Alzheimer’s Disease (AD) and are classified as *APOEε2, APOEε3*, and *APOEε4*. The protein products of these three alleles differ from each other by single amino-acid substitutions at positions 112 and 158 [1]. This leads to conformational changes in two alpha helices and causes the APOEε4 protein to be more compact, less protected and less stable compared to APOEε2 and APOEε3 [1]. Furthermore, these changes lead to functional differences in lipid binding [1]. The *APOEε4* allele increases the risk of AD relative to *APOEε3*, while the *APOEε2* allele decreases risk [2], [3]. *APOEε4* is also associated with earlier age of AD onset by approximate 2.5 years [2]. The amyloid cascade hypothesis and the tau hypothesis are the two most commonly-accepted hypotheses of AD pathology [4], [5]. APOE is potentially involved in both hypotheses. In addition to altering lipid binding, APOE potentially plays an important role in amyloid-β (Aβ) protein deposition [4], [6]. Human studies show that the *APOEε4* allele dosage is associated with increased Aβ plaques in AD patients [7], and *APOEε4* carriers who are middle-aged or elderly are more likely to have brain amyloid while *APOEε2* carriers rarely develop fibrillar Aβ [8]–[10]. *In vitro* experiments also show that the APOE protein binds to Aβ with high avidity [11], and mouse model experiments suggest that APOE regulates Aβ metabolism, aggregation, and deposition [12], [13]. APOE also clears soluble Aβ in mice, with *APOEε4* less efficient than *APOEε2* or *APOEε3* [14]. In addition, APOE affects tau neuropathological changes in AD brains [15]. Abnormal tau phosphorylation was found in apoE4 mice brains [16].

The lipid clearance function of APOE is also influenced by the oxidative state of the protein, with oxidized APOE showing lower lipid-binding affinity [17]. Haptoglobin (HP) is a hemoglobin scavenger that keeps free hemoglobin from causing oxidative damage to tissues, and HP is also a potential antioxidant of APOE. *In vitro* experiments have shown that the HP protein physically binds to APOE and this binding potentially protects the APOE protein against oxidation and preserves its lipid transport activity [18], [19].

A common structural variant (SV) of the *HP* gene spans two tandem exons and distinguishes two alleles: *HP1* (one copy of exons 3 and 4) and *HP2* (two copies of exons 3 and 4) [20]. This variant is not captured via genotyping arrays that are typically used in genome-wide association studies (GWAS) due to the complexity of the surrounding linkage disequilibrium and haplotype structures, so the effects of this variant have not been adequately explored in prior large-scale GWAS studies. This SV alters the α-subunit of HP, changing the quaternary structure of the final HP protein complex [21]. HP1 proteins form dimers, the HP1 and HP2 together form multimers with linear conformations, and HP2 forms multimers with cyclic conformations [22], [23]. Though the hemoglobin binding affinity of the HP protein complex is not strongly influenced, due to their larger sizes, multimers demonstrate reduced binding capability, thereby leading to lower functional activity compared to dimers [22]. This SV has been previously associated with plasma lipid levels, and (in our prior work) with neurocognitive deficits in people living with HIV [20], [24], [25]. A small study also reported that the interaction of *APOEε4* and *HP* genotypes associated with longevity in a population in central Italy [26].

Given the antioxidant role of HP, previously reported associations to cognitive phenotypes, and its physical interactions with APOE, here we investigate the genetic interaction between *HP* and *APOE* polymorphisms and its effect on AD risk and age of onset in the largest collection of AD samples in the U.S.

## 2 METHODS

### 2.1 Study Cohorts and Phenotype

The study data includes European-descent participants from 29 Alzheimer’s Disease Genetic Consortium (ADGC) cohorts with available genotyping data (Supplemental Text 1). The sample size and other descriptive statistics are provided in **Table 1**. The detailed description of each cohort along with their diagnosis of AD can be found at [27]. Age at AD onset is available for a subset of cohorts including the ADCs, TGEN2, NIA-LOAD, MIRAGE, ACT, UPITT, and ROS/MAP [28]. A detailed description of the ascertainment of age at AD onset along with the descriptive statistics of each cohort can be found at [29]. All subjects were recruited under protocols approved by the appropriate Institutional Review Boards. Written informed consent was obtained from study participants or, for those with substantial cognitive impairment, from a caregiver, legal guardian, or other proxy.

**Table 1.** Study Population

| Overall |  | Per Cohort |  |  |
| --- | --- | --- | --- | --- |
| N | 24102 | Cohort | N (%) | *HP* Hard-call Rate (%) |
| Categorical Variables [N (%)] |  | ACT1 | 2036 (8.4) | 96.94 |
| *HP* Genotype |  | ACT2 | 29 (0.1) | 96.87 |
| *HP1/HP1* | 3494 (14.5) | ADC1 | 2001 (8.3) | 96.72 |
| *HP1/HP2* | 11157 (46.3) | ADC2 | 854 (3.5) | 96.98 |
| *HP2/HP2* | 9451 (39.2) | ADC3 | 1370 (5.7) | 97.28 |
| *APOE* Genotype |  | ADC4 | 659 (2.7) | 97.06 |
| $\epsilon 33$ | 10928 (47.3) | ADC5 | 769 (3.2) | 97.3 |
| $\epsilon 22$ | 88 (0.4) | ADC6 | 526 (2.2) | 97.15 |
| $\epsilon 23$ | 1881 (8.1) | ADC7 | 1269 (5.3) | 97.68 |
| $\epsilon 24$ | 578 (2.5) | ADNI | 425 (1.8) | 96.68 |
| $\epsilon 34$ | 7822 (33.9) | BIOCARD | 115 (0.5) | 98.02 |
| $\epsilon 44$ | 1796 (7.8) | CHAP2 | 167 (0.7) | 97.44 |
| Sex |  | EAS | 146 (0.6) | 97.54 |
| Female | 14241 (59.1) | GSK | 1135 (4.7) | 82.19 |
| Male | 9861 (40.9) | NIA-LOAD | 3250 (13.5) | 96.85 |
| AD |  | MAYO | 1665 (6.9) | 97.85 |
| Case | 11684 (48.5) | MIRAGE | 1189 (4.9) | 98.07 |
| Control | 12418 (51.5) | NBB | 125 (0.5) | 97 |
| Numerical Variables [mean (SD)] |  | OHSU | 274 (1.1) | 96.71 |
| Age at AD onset | 74.18 (7.68) | RMAYO | 239 (1.0) | 97.45 |
| Age at last visit | 77.77 (8.12) | ROSMAP1 | 1007 (4.2) | 96.97 |
| Age combined | 75.45 (8.03) | ROSMAP2 | 268 (1.1) | 95.87 |
|  |  | TARCC | 456 (1.9) | 91.68 |
|  |  | TGEN2 | 761 (3.2) | 72.93 |
|  |  | UKS | 742 (3.1) | 97.31 |
|  |  | UPITT | 1375 (5.7) | 66.74 |
|  |  | WASHU | 512 (2.1) | 97.61 |
|  |  | WASHU2 | 128 (0.5) | 97.45 |
|  |  | WHICAP | 610 (2.5) | 96.91 |

The “N”s are the number of individuals that are included in the analyses after omitting for missing in AD status. The “Age combined” variable is equal age at onset for AD cases and age at last visit for controls.

### 2.2 Genotyping and QC

Genotyping was performed on either Illumina or Affymetrix high-density SNP microarrays. For Illumina chip data, a minimal call rate of 0.95 and a minor allele frequency (MAF) of 0.02 were used for filtering and for Affymetrix 0.98 and 0.01 were used, respectively [30].

Ancestry-based principal components (PCs) were computed from a combined dataset using the set of SNPs genotyped in all study cohorts [28]. After filtering SNPs with pairwise LD (r2) <0.20, 31,310 SNPs were evaluated using EIGENSTRAT [28]. The top three PCs from EIGENSTRAT were used as covariates in the joint analysis [28].

The *APOE* genotype was determined for different cohorts in multiple ways including using SNPs rs7412 and rs429358, Roche Diagnostics LightCycler 480 (Roche Diagnostics) instrument LightMix Kit ApoE C112R R158 (TIBMOLBIOL), pyrosequencing or restriction fragment length polymorphism analysis, and high-throughput sequencing of codons 112 and 158 in *APOE* by Agencourt Bioscience Corporation [28].

### 2.3 Imputation

After QC, SNP data for chromosome 16 was extracted from available genotype data, submitted to the TOPMed imputation server [31]–[33], and imputed to the TOPMed (version TOPMed-r2) genotype marker set for each cohort. This approach was validated using gene-tissue expression (GTEx) RNA expression data (Supplemental Text 2, Table S1, S2, Figure S1). To impute the *HP* SV, we used a customized version of a published imputation reference that was developed using droplet PCR and validated using RNA-sequencing [20], [25], [Supplement File --Imputation Reference]. After TOPMed imputation, we first extracted SNPs that are included in the HP imputation reference. We then filtered the SNPs for TOPMed imputation R^2^ value and kept only markers that had an R^2^ ≥0.8. We performed HP imputation with the customized reference panel using IMPUTEv2 software for each cohort [34]. Finally, we conducted hard-calling for the imputed *HP* allele dosages with a threshold of 0.9. Only imputation dosages ≥0.9 were kept for further analyses and dosages <0.9 were removed. This approach has been previously validated [25].

### 2.4 Association Analysis

Association analyses were conducted jointly using data from all cohorts together. We sought to model the effect of *APOE* alleles comprehensively, so we combined both the detrimental effect of *APOEε4* and the protective effect of *APOEε2*. To accomplish this, we assumed equal change in the AD risk between *APOEε2* to *APOEε3*, and *APOEε3* to *APOEε4* alleles in our statistical analyses, and model each allele separately (referred below as “*APOE1ε2-3-4*” and “*APOE2ε2-3-4*”). Logistic regression models with AD case/control status as an outcome were fit for all individuals with an interaction term of *HP2* allele count and *APOE* allele effects.

All regression analyses were adjusted for sex, age, and the top 3 ancestry-based PCs. Age is defined as the age at onset for AD cases and age at last visit for controls. All analyses were conducted using R (Metafor [35], Survival [36], [37]). The statistical models used in analyses are as below:

We fit a logistic regression model using the full data with an *APOEε2-3-4* effect for each of the two *APOE* alleles:

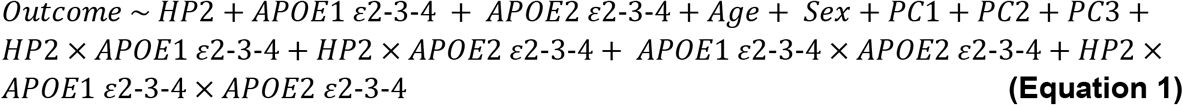

where the “*HP2*” represents the number of *HP2* alleles, and “*APOE1 ε2-3-4”* and

*“APOE2 ε2-3-4”* represents the individual “*APOEε2-3-4*” effect of each *APOE* allele.

We also fit individual stratified logistic regression models for the three *APOE* strata (i.e. individuals who carry at least one *ε2, ε3*, or *ε4* allele, respectively) as shown below:

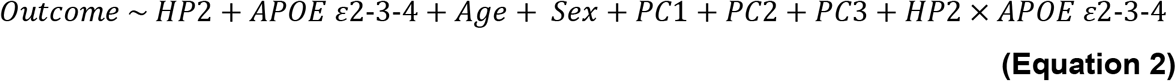

where the “*APOE ε2-3-4*” represents the “*APOEε2-3-4*” effect of the remaining *APOE* allele within each stratus.

In addition, we conducted survival analysis using the Cox proportional hazards model to investigate the effects of *HP* on the age of AD onset. The model is as below:

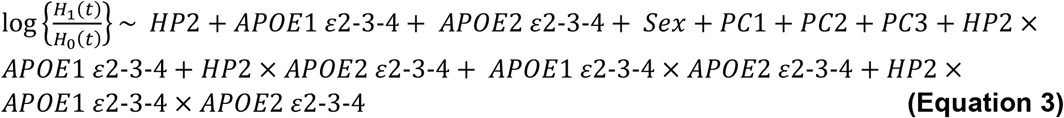

where the “*H*_*i*_*(t)*” represents the hazard. The sample size (N) for analyses may differ due to variable missingness within subsets.

## 3 RESULTS

To examine the *HP* SV for association to AD risk, we imputed the *HP* SV for European-descent ADGC participants across 29 cohorts originally genotyped with multiple different arrays (**Table 1**). We obtained a >95% hard-call rate in 25/29 of the cohorts, with an overall hard-call rate of 93.52% (33,725/36,062). This call rate suggests high imputation quality and confidence despite heterogeneity in genotyping platform. We noted that 4/29 cohorts showed statistically significant deviations from Hardy-Weinberg Equilibrium; however, sensitivity analyses removing these cohorts showed no qualitatively different results.

We first studied the association between AD status and *HP2* allele count adjusting for sex, age, and the top three ancestry-based PCs. No statistically significant effect of *HP* genotypes was found (p=0.23). We also examined dominant and recessive models of the *HP2* alleles and found no significant associations (p=0.26 and p=0.37, respectively).

Given prior evidence of their molecular interaction, we further investigated *HP* alleles in the context of *APOE*. In these models, for simplification we assumed an equal (linear) increase in AD risk from *APOEε2* to *APOEε3*, and *APOEε3* to *APOEε4* (denoted as *APOEε2-3-4*) (see Supplemental Text 3 for more details on how we arrived at our modeling strategy). We first fit a logistic regression model using AD status as the outcome with age, sex, the first 3 PCs, *HP2* allele count, *APOEε2-3-4* for each of the two *APOE* alleles, and pairwise interactions and a three-way interaction of these genetic effects of the two *APOE* alleles and the *HP* genotype (**Equation 1**). We found significant main effects from each of the two *APOEε2-3-4* variables and *HP2* allele count, along with their two-way and three-way interaction terms (**Table 2**). These interaction effects can be observed in **Figure 1A**, whereby AD risk decreases dramatically with each *HP2* allele in *APOEε24* (purple) individuals, yet increases in *APOEε44* (dark green), *APOEε23* (light green) and *APOEε22* (yellow) individuals. The risk remained fairly equal in *APOEε33* (magenta) and *APOEε34* (orange) individuals regardless of *HP2* allele count (**Figure 1A**).

**Table 2.** Effect of *HP* and both *APOE* alleles on AD risk (N=22,651).

| Variable | OR | 95%CI | p_value |
| --- | --- | --- | --- |
| (Intercept) | 0.03 | (0.01, 0.12) | 2.372e-06 |
| ***HP2*** | **3.99** | **(1.43, 11.09)** | **0.008** |
| ***APOE1* ε2-3-4** | **2.89** | **(1.39, 5.99)** | **0.004** |
| ***APOE2* ε2-3-4** | **4.76** | **(2.70, 8.39)** | **6.891e-08** |
| Sex: Female | 0.98 | (0.93, 1.04) | 0.547 |
| Age | 0.98 | (0.98, 0.98) | 3.267e-27 |
| PC1 | 0.69 | (0.28, 1.68) | 0.410 |
| PC2 | 0.45 | (0.18, 1.10) | 0.078 |
| PC3 | 1.61 | (0.64, 4.03) | 0.308 |
| ***HP2* x *APOE1* ε2-3-4** | **0.49** | **(0.29, 0.82)** | **0.007** |
| ***HP2* x *APOE2* ε2-3-4** | **0.56** | **(0.38, 0.84)** | **0.005** |
| *APOE1* ε2-3-4 x *APOE2* ε2-3-4 | 0.85 | (0.64, 1.12) | 0.236 |
| ***HP2* x *APOE1* ε2-3-4 x *APOE2* ε2-3-4** | **1.34** | **(1.10, 1.63)** | **0.004** |

Effects are from a logistic regression model described in **Equation 1**. *HP2* was included as the count of *HP2* alleles. “*APOE1* ε2-3-4” and “*APOE2* ε2-3-4” represents the individual *APOE*ε2-3-4 effects from the two copies of *APOE* alleles, respectively. “Age” represents the age at onset for AD cases and age at last visit for controls. N is the number of individuals in this model after omitting missing values in all model variables.

**Figure 1.**
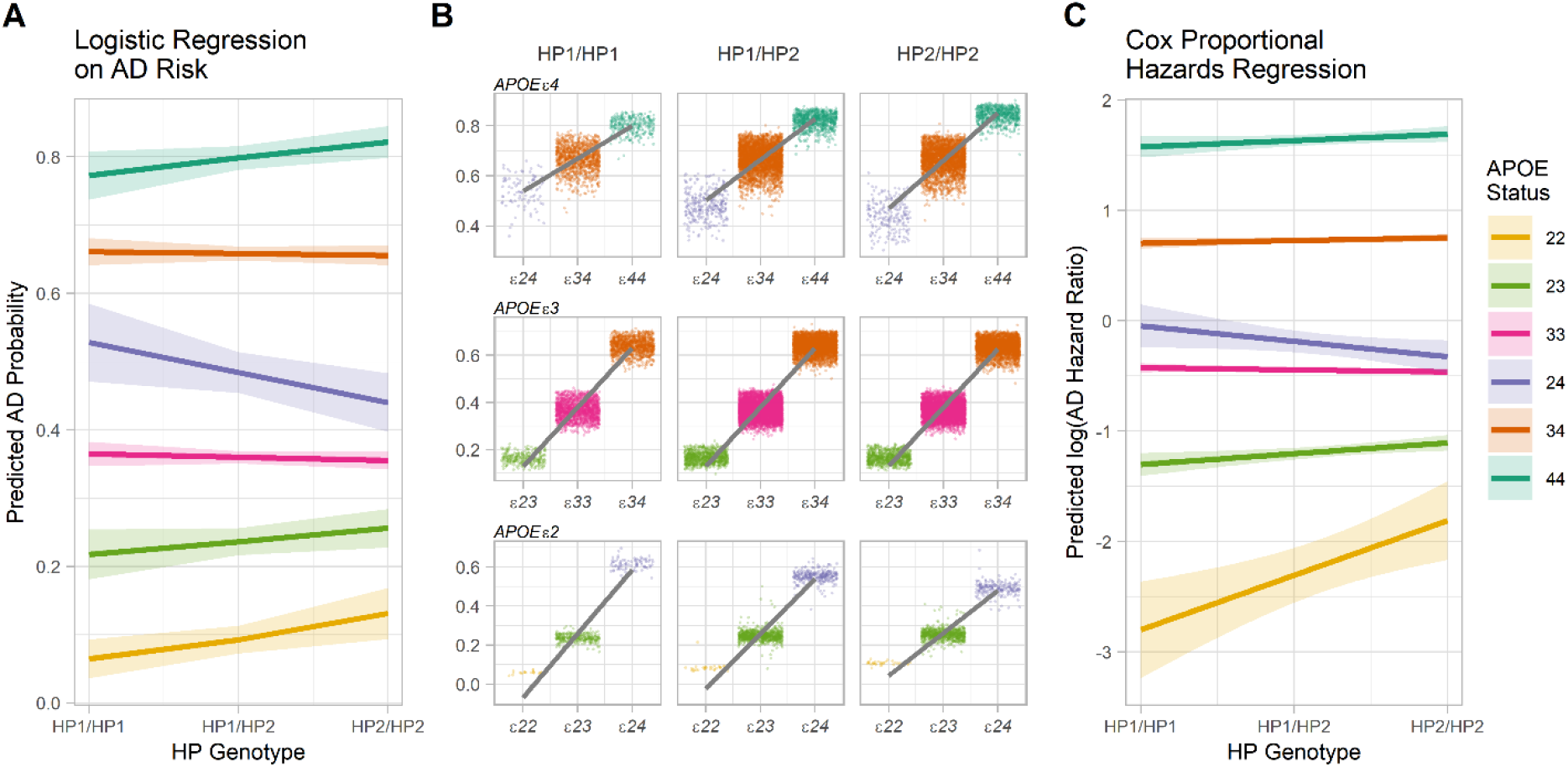
Effect of *APOE* and *HP* Genotype on AD risk and Age. **(A)** Trend lines show the fitted effect estimates and standard errors from the logistic regression model in **Table 2. (B)** Jitter plot and trend lines show the linear relationship between *APOEε2-3-4* and predicted AD probability stratified by *HP* genotypes for all *APOE* strata. Note the increase in slope of the trendline with each additional *HP2* allele for *ε4* carriers (left to right of the top row) and the decrease in slope of the trendline with each additional *HP2* allele for *ε2* carriers (left to right of the bottom row) **(C)** Trend lines show the fitted effects estimates and standard errors from the Cox proportional hazards regression model in **Table 4**.

To more easily describe the interaction effects detected in this model, we also fit individual logistic regression models stratified by participants’ *APOE* genotype, as illustrated in **Equation 2**. This approach isolates the effect of one *APOE* allele on the background of *ε2, ε3*, or *ε4*. Among *APOEε4* carriers (i.e. people with *APOEε24, ε34*, and *ε44*, N=9,949), the *APOEε2-3-4* effect of the remaining allele leads to increased AD risk as expected (p=5.477e-8, **Table 3**). Each *HP2* allele further increases this *APOEε2-3-4* effect on AD risk significantly (p=0.025, **Table 3, Figure 1B Top**). This can be seen as an increasing slope in the trend of the average risk (seen as increasing steepness of the grey trend lines in **Figure 1B Top** from left to right) with each additional *HP2* allele. Among *APOEε2* carriers (n=2,515), given one *APOEε2* allele, the *APOEε2-3-4* effect of the remaining allele also leads to an increased AD risk (p=1.689e-16, **Table 3**). However, unlike the *APOEε4* carriers, each *HP2* allele decreases this *APOEε2-3-4* effect on AD risk (p=0.035, **Table 3, Figure 1B Bottom**). This can be seen as decreasing slope in the trend of the average risk (seen as decreasing steepness of the grey trend lines in **Figure 1B Bottom** from left to right) with each additional *HP2* allele. Analyses of *APOEε3* carriers did not show any significant effects of the *HP* alleles or any significant *HP-APOE* interactions (**Table 3, Figure 1B Middle**). Thus, the significant interactions in our model are due to *APOEε4* and *APOEε2* carriers, where the *HP* alleles show opposite modifying effects of *APOE* on AD risk.

**Table 3.**
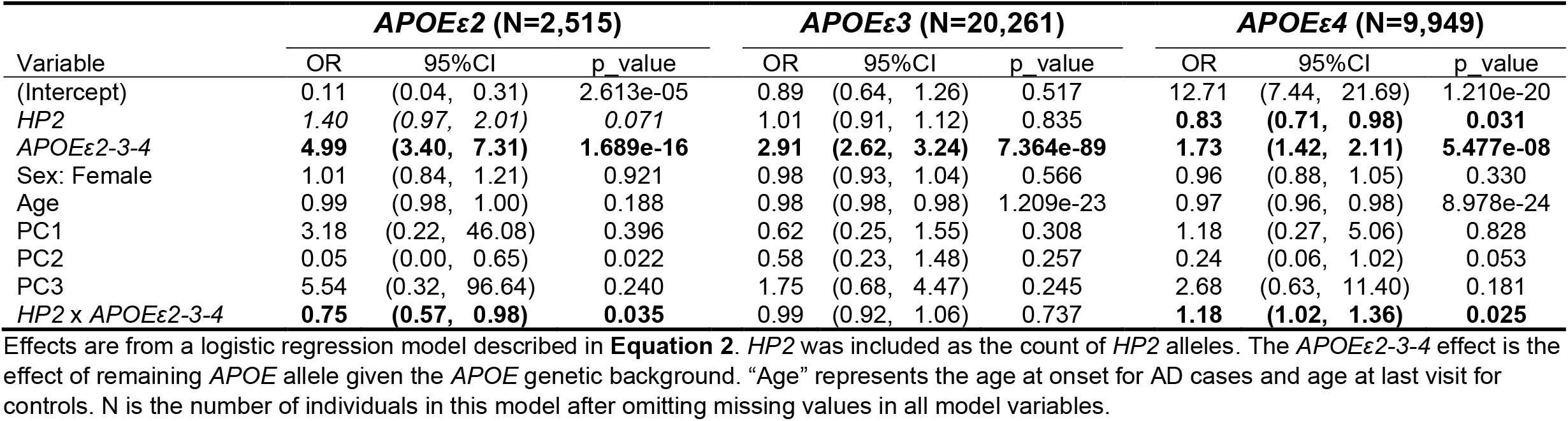
Effect of *HP* and *APOEε2-3-4* alleles on AD risk stratified by *APOE* genetic background.

| Variable | *APOE*ε2 (N=2,515) | | | *APOE*ε3 (N=20,261) | | | *APOE*ε4 (N=9,949) | | |
| --- | --- | --- | --- | --- | --- | --- | --- | --- | --- |
|  | OR | 95%CI | p_value | OR | 95%CI | p_value | OR | 95%CI | p_value |
| (Intercept) | 0.11 | (0.04, 0.31) | 2.613e-05 | 0.89 | (0.64, 1.26) | 0.517 | 12.71 | (7.44, 21.69) | 1.210e-20 |
| *HP*2 | 1.40 | (0.97, 2.01) | 0.071 | 1.01 | (0.91, 1.12) | 0.835 | **0.83** | **(0.71, 0.98)** | **0.031** |
| *APOE*ε2-3-4 | **4.99** | **(3.40, 7.31)** | **1.689e-16** | **2.91** | **(2.62, 3.24)** | **7.364e-89** | **1.73** | **(1.42, 2.11)** | **5.477e-08** |
| Sex: Female | 1.01 | (0.84, 1.21) | 0.921 | 0.98 | (0.93, 1.04) | 0.566 | 0.96 | (0.88, 1.05) | 0.330 |
| Age | 0.99 | (0.98, 1.00) | 0.188 | 0.98 | (0.98, 0.98) | 1.209e-23 | 0.97 | (0.96, 0.98) | 8.978e-24 |
| PC1 | 3.18 | (0.22, 46.08) | 0.396 | 0.62 | (0.25, 1.55) | 0.308 | 1.18 | (0.27, 5.06) | 0.828 |
| PC2 | 0.05 | (0.00, 0.65) | 0.022 | 0.58 | (0.23, 1.48) | 0.257 | 0.24 | (0.06, 1.02) | 0.053 |
| PC3 | 5.54 | (0.32, 96.64) | 0.240 | 1.75 | (0.68, 4.47) | 0.245 | 2.68 | (0.63, 11.40) | 0.181 |
| *HP*2 x *APOE*ε2-3-4 | **0.75** | **(0.57, 0.98)** | **0.035** | 0.99 | (0.92, 1.06) | 0.737 | **1.18** | **(1.02, 1.36)** | **0.025** |

Effects are from a logistic regression model described in **Equation 2**. *HP*2 was included as the count of *HP*2 alleles. The *APOE* ε2-3-4 effect is the effect of remaining *APOE* allele given the *APOE* genetic background. “Age” represents the age at onset for AD cases and age at last visit for controls. N is the number of individuals in this model after omitting missing values in all model variables.

**Table 4.**
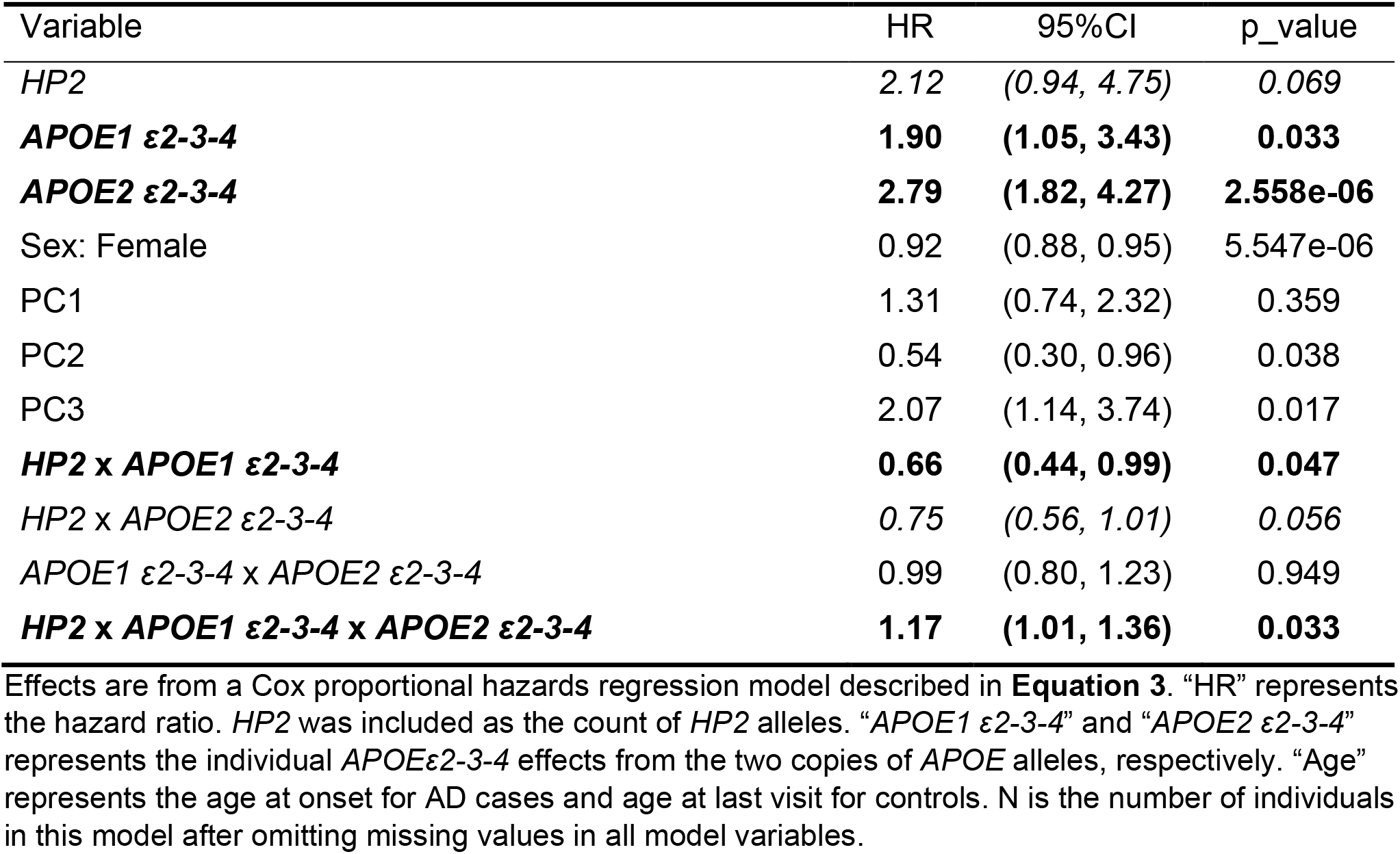
Effect of *HP* and both *APOE* alleles on age at AD onset (N=22,651).

| Variable | HR | 95%CI | p_value |
| --- | --- | --- | --- |
| *HP2* | 2.12 | (0.94, 4.75) | 0.069 |
| ***APOE1* ε2-3-4** | **1.90** | **(1.05, 3.43)** | **0.033** |
| ***APOE2* ε2-3-4** | **2.79** | **(1.82, 4.27)** | **2.558e-06** |
| Sex: Female | 0.92 | (0.88, 0.95) | 5.547e-06 |
| PC1 | 1.31 | (0.74, 2.32) | 0.359 |
| PC2 | 0.54 | (0.30, 0.96) | 0.038 |
| PC3 | 2.07 | (1.14, 3.74) | 0.017 |
| ***HP2* x *APOE1* ε2-3-4** | **0.66** | **(0.44, 0.99)** | **0.047** |
| *HP2* x *APOE2* ε2-3-4 | 0.75 | (0.56, 1.01) | 0.056 |
| *APOE1* ε2-3-4 x *APOE2* ε2-3-4 | 0.99 | (0.80, 1.23) | 0.949 |
| ***HP2* x *APOE1* ε2-3-4 x *APOE2* ε2-3-4** | **1.17** | **(1.01, 1.36)** | **0.033** |

Effects are from a Cox proportional hazards regression model described in **Equation 3**. “HR” represents the hazard ratio. *HP2* was included as the count of *HP2* alleles. “*APOE1* ε2-3-4” and “*APOE2* ε2-3-4” represents the individual *APOE* ε2-3-4 effects from the two copies of *APOE* alleles, respectively. “Age” represents the age at onset for AD cases and age at last visit for controls. N is the number of individuals in this model after omitting missing values in all model variables.

We have previously shown that genetic interaction models can show false positive associations due to deviation from the model’s genotype effect assumption [38], which in our case is congruent to the dose-response assumption for *APOEε2-3-4*. Therefore, we further explored the impact of our linear *APOEε2-3-4* assumption on the model fitting. While this simplifying assumption lets us assess the change in *APOE* effect due to the *HP* alleles, prior literature suggests that the protective effect of *ε2* is smaller in magnitude than the risk effect of *ε4*. We conducted a sensitivity analysis exploring the individual effect of *ε3* and the results demonstrated that the *HP* effect of *APOE* on AD risk is not due to misspecification of *APOE* main effects (Supplemental Text 3, Table S3 – S5).

Given the known association of *APOE* to age of AD onset, we hypothesized that the *HP-APOE* interactions might also influence/modify the *APOE* effect on the age of AD onset. Thus, we performed survival analysis using a Cox proportional hazards regression model to investigate the effects of *APOEε2-3-4* from both *APOE* alleles individually and *HP2* allele count on the age at AD onset (**Equation 3**) with controls included as censored. Similar significant effects were found for our age of onset analysis as were found for AD risk, though some coefficients were of marginal significance (**Table 4, Figure 1C**). The directions of these effects (positive/negative) were congruent to our findings in logistic regression models.

An eQTL (expression quantitative trait locus) — rs2000999 is associated with the RNA level of *HP* that is independent of the *HP* SV [19][20]. We evaluated the impact of the genotypes of this eQTL and found no impact on our results (Supplemental Text 4, Table S6).

As prior work suggests divergent evolutionary histories of this *HP* variant and potentially different effects of *HP* alleles on neurocognition by ancestry [20], we also evaluated the *HP-APOE* interaction in African American (AA) ADGC cohorts. We imputed the *HP* genotypes for 12 ADGC AA cohorts (N=4,617, Supplemental Text 5). Although we are underpowered to replicate the *HP-APOE* interaction effect in the AA cohorts (Table S7), a random effects meta-analysis for both European and AA cohorts’ data showed that the detected associations had consistent directions of effect and retained statistical significance (**Figure 2**, Supplemental Text 6). Also, both the main effect of *HP2* and the interaction effect of *HP2* and *APOEε2-3-4* became more significant in *APOEε2* individuals (Table S8, S9) and less significant in *APOEε4* individuals, which is consistent with a previous report that *APOEε4* confers a lower risk to AD in AA individuals [39].

**Figure 2.**
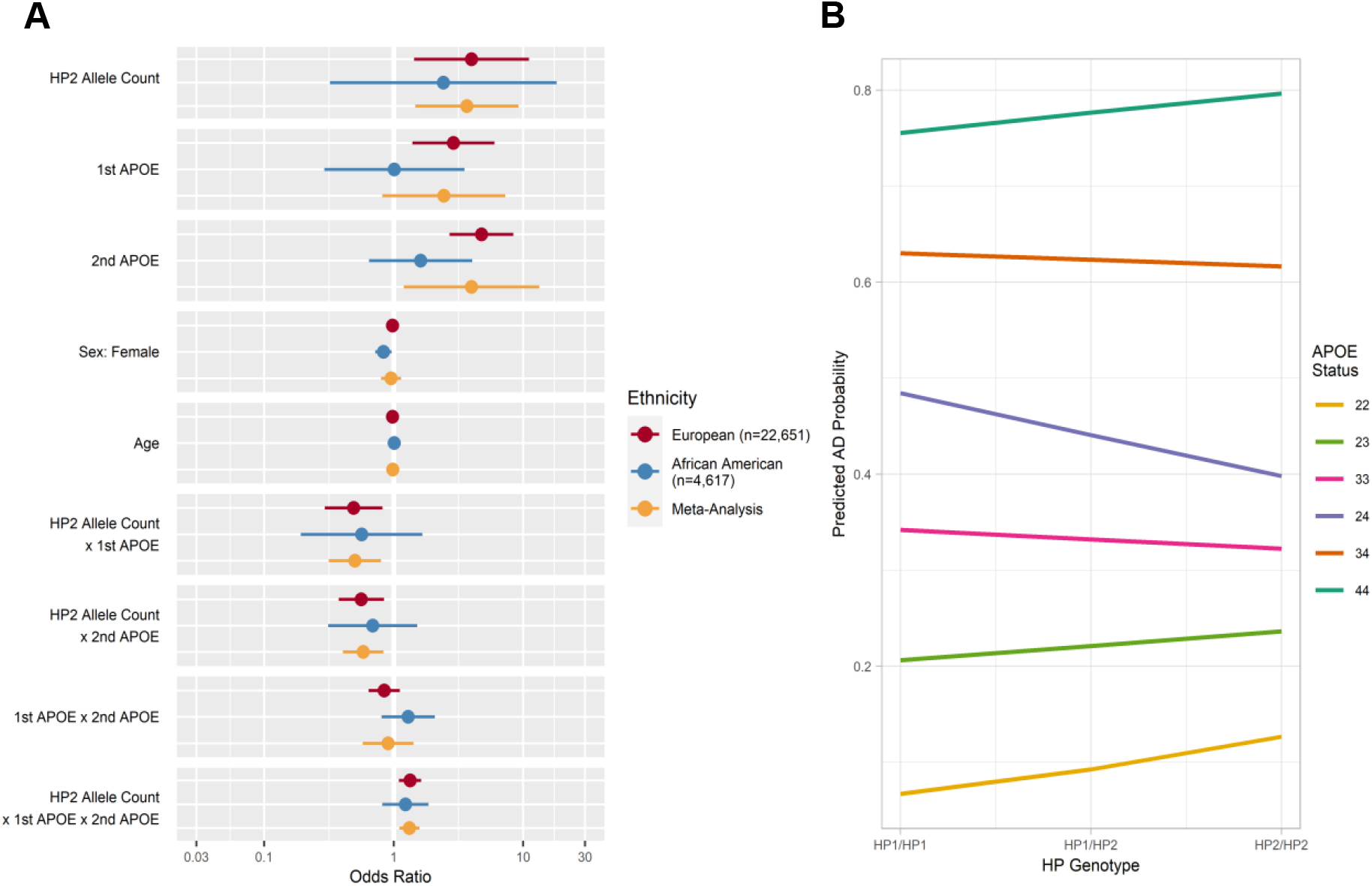
Random Effects Meta-Analysis of European and AA Data. European population effects are from the logistic regression model shown in **Table 2**. AA population effects are from the logistic regression model shown in Table S7. Meta-analysis effects are from a random effects meta-analysis of both European and African American data shown in Table S8.

## 4. STRENGTHS AND LIMITATIONS

The ADGC dataset provided a large sample size to statistically evaluate this important HP-APOE molecular hypothesis, and we have achieved a high accuracy in our validated SV imputation process. The strength of our reported effect is also bolstered by a meta-analysis across different ancestry groups, and is strongly supported by prior mechanistic evidence of HP protein interactions with known components of AD risk.

This study design and analysis has limitations. All the study subjects have an age of >60, thus we can make no inference on the role of HP in early onset forms of AD. As with prior studies of ADGC cohorts, we noticed large heterogeneity in age and case/control distributions across cohorts due to differences in geography and ascertainment strategy. This heterogeneity did not impact HP imputation; however, this could impact individual analyses within each cohort. The *HP* genotypes from 4 out of 29 cohorts are significantly deviated from Hardy-Weinberg Equilibrium (HWE). However, similar effects were replicated in the 25 cohorts with no qualitative change in results suggesting that the effects were not driven by this deviation. We and others have also reported associations between the *HP* SV and serum and CSF haptoglobin levels where the genetic effect of the *HP* SV remains independent of the CSF HP levels. Due to the limited availability of this data in our AD cohorts, we cannot eliminate haptoglobin levels as a potential confounder of our association.

## 5. DISCUSSION

The epsilon alleles of the *APOE* gene have been repeatedly shown to impact AD risk in a dose-dependent fashion. As a result, many studies adjust for the *APOE* effect by enumerating the number of *ε4* alleles and often ignoring the more infrequent protective *ε2* alleles. Here, we show that a functional SV in the *HP* gene alters the effect of *APOE* alleles on AD risk in ways that defy this convention. For example, the *APOEε24* genotype is generally considered to have higher risk for AD given the detrimental effect of *ε4*. However, individuals with *APOEε24* and *HP2/HP2* are closer in risk to *APOEε33* individuals than *APOEε34* or *APOEε44* individuals. These results suggest that for some scenarios where APOE stratification or adjustment is needed, the inclusion of *HP* genotypes and interactions is likely to improve predictions of AD risk.

There are multiple potential biological hypotheses that could drive the statistical interaction we detected. Both APOE and HP are related to inflammation. *In vitro* experiments using human *APOE* knock-in mice showed that *APOEε4* mice are more susceptible to inflammation compared to *APOEε2* and *APOEε3* [40]. The APOE peptide inhibits inflammation processes in isolated microglia [40]; furthermore, microglia with an *APOEε4* background demonstrated a greater release of pro-inflammatory cytokines [41]. HP, as a free hemoglobin scavenger, reduces oxidative stress by preventing the release of free heme iron, thereby reducing inflammation, generation of reactive oxygen species, and oxidative damage to surrounding tissues. It is hypothesized that *HP* alleles exhibit differing antioxidant activities; therefore the production of and protection from inflammatory products could be balanced by the differential activities of the *APOE* and *HP* alleles in a way that produces the interaction effect we observed. Studies have also shown associations between *HP* alleles and vascular complications of diabetes [42], which may point to a role for this interaction in the vascular contributions to dementia. More directly, HP physically binds to and is an antioxidant of the APOE protein, likely protecting APOE function and activity. HP2 offers lower oxidation protection compared to HP1. However, due to the conformational differences in APOE structure from allelic variants, the HP antioxidant activity may also depend on the *APOE* allele. If true, this hypothesis would explain why we observe a statistical interaction effect of *HP* and *APOE* on AD risk rather than an independent effect of *HP*.

HP may also play a role in the binding of APOE with Aβ. Spagnuolo *et al* found that HP promotes formation of stable Aβ complexes with APOE proteins. Immunoassays showed that HP binds to Aβ in brain tissue from AD patients [43]. Shi *et al* found that APOE and HP, along with 4 more proteins, are consistently associated with high amyloid burden [44]. Esiri *et al* showed that HP facilitates the binding of APOE and Aβ [45]. It was also reported that in the human glioblastoma–astrocytoma cell line U-87 MG, HP impairs Aβ uptake and limits the toxicity of this peptide on these cells [46]. Therefore, another mechanistic hypothesis from our findings could be that these *HP* alleles may alter the promotion of binding of Aβ to APOE in an *APOE*-allele specific manner.

This variant affecting HP function may alter strategies for using APOE as a therapeutic target for AD [47], or efforts to modify APOE-Aβ binding [48]. Haptoglobin has been used as a therapeutic agent in some settings for over three decades and has also been explored as way to mitigate oxidative damage from hemoglobin-driven pathology in the brain [49]. Given its prior clinical use and the ability to synthesize both HP functional alleles [50], a form of haptoglobin therapy could potentially be tailored to individuals based on *APOE* genotype.

## ACKNOWLEDGEMENTS

The NACC database is funded by NIA/NIH Grant U01 AG016976. NACC data are contributed by the NIA-funded ADCs: P30 AG019610 (PI Eric Reiman, MD), P30 AG013846 (PI Neil Kowall, MD), P50 AG008702 (PI Scott Small, MD), P50 AG025688 (PI Allan Levey, MD, PhD), P30 AG010133 (PI Andrew Saykin, PsyD), P50 AG005146 (PI Marilyn Albert, PhD), P50 AG005134 (PI Bradley Hyman, MD, PhD), P50 AG016574 (PI Ronald Petersen, MD, PhD), P50 AG005138 (PI Mary Sano, PhD), P30 AG008051 (PI Steven Ferris, PhD), P30 AG013854 (PI M. Marsel Mesulam, MD), P30 AG008017 (PI Jeffrey Kaye, MD), P30 AG010161 (PI David Bennett, MD), P30 AG010129 (PI Charles DeCarli, MD), P50 AG016573 (PI Frank LaFerla, PhD), P50 AG016570 (PI David Teplow, PhD), P50 AG005131 (PI Douglas Galasko, MD), P50 AG023501 (PI Bruce Miller, MD), P30 AG035982 (PI Russell Swerdlow, MD), P30 AG028383 (PI Linda Van Eldik, PhD), P30 AG010124 (PI John Trojanowski, MD, PhD), P50 AG005133 (PI Oscar Lopez, MD), P50 AG005142 (PI Helena Chui, MD), P30 AG012300 (PI Roger Rosenberg, MD), P50 AG005136 (PI Thomas Grabowski, MD, PhD), P50 AG033514 (PI Sanjay Asthana, MD, FRCP), and P50 AG005681 (PI John Morris, MD).

Samples from the National Cell Repository for Alzheimer’s Disease (NCRAD), which receives government support under a cooperative agreement grant (U24 AG21886) awarded by the National Institute on Aging (NIA), were used in this study. We thank contributors who collected samples used in this study, as well as patients and their families, whose help and participation made this work possible; Data for this study were prepared, archived, and distributed by the National Institute on Aging Alzheimer’s Disease Data Storage Site (NIAGADS) at the University of Pennsylvania (U24-AG041689–01).

The funders had no role in study design, data collection and analysis, decision to publish, or preparation of the manuscript.

Trans-Omics in Precision Medicine (TOPMed) program imputation panel (version TOPMed-r2) supported by the National Heart, Lung and Blood Institute (NHLBI); see www.nhlbiwgs.org. TOPMed study investigators contributed data to the reference panel, which can be accessed through the Michigan Imputation Server; see https://imputationserver.sph.umich.edu. The panel was constructed and implemented by the TOPMed Informatics Research Center at the University of Michigan (3R01HL-117626-02S1; contract HHSN268201800002I). The TOPMed Data Coordinating Center (3R01HL-120393-02S1; contract HHSN268201800001I) provided additional data management, sample identity checks, and overall program coordination and support.

We gratefully acknowledge the studies and participants who provided biological samples and data for TOPMed.

We also acknowledge the Genotype-Tissue Expression Project. The Genotype-Tissue Expression Project was supported by the Common Fund of the Office of the Director of the National Institutes of Health, and by NCI, NHGRI, NHLBI, NIDA, NIMH, and NINDS. The data used for the analyses described in this manuscript were obtained from the GTEx Portal on 03/08/2018 and dbGaP accession number phs000424.vN.pN on 03/08/2018.

