## Supplement for "A Haptoglobin (HP) Structural Variant Alters the Effect of *APOE* Alleles on Alzheimer’s Disease"

### Supplement Material for: A Haptoglobin (HP) Exon Deletion Polymorphism Alters the Effect of APOE Alleles on Alzheimer's Disease

#### Supplement Text:

##### 1. Study Cohorts

The Alzheimer's Disease Genetic Consortium (ADGC) is composed of 31 cohorts including the National Institute on Aging (NIA) Alzheimer's Disease Centers (ADCs), the Adult Changes in Thought (ACT) study [1], the Alzheimer Disease Neuroimaging Initiative (ADNI) study [2], the National Institute on Aging Late-Onset Alzheimer's Disease Family (NIA-LOAD) study [3], the Mayo Clinic Jacksonville (MAYO), the Multi Institutional Research of Alzheimer Genetic Epidemiology (MIRAGE) study [4], Oregon Health and Science University (OHSU), the Rush University Religious Orders Study/Memory and Aging Project (ROSMAP) [5], the Texas Alzheimer's Research and Care Consortium (TARCC), the Translational Genomics Research Institute series 2 (TGEN2) [6], University of Pittsburgh (UPitt) [7], Washington University (WASHU), the Universitätsklinikum des Saarlandes (UKS), and the Netherlands Brain Bank (NBB) [8], Biomarkers of Cognitive Decline Among Normal Individuals: the BIOCARD cohort (BIOCARD), Chicago Health and Aging Project (CHAP2), Einstein Aging Study (EAS), Mayo Clinic (RMAYO), Washington Heights-Inwood Community Aging Project (WHICAP) [9]. Detailed inclusion and exclusion criteria can be found from previous publications [8]–[10].

##### 2. TOPMed Background Imputation Validation

The HP imputation marker coverage of directly genotyped ADGC data was not high. After QC, the majority of the cohorts had a coverage less than 50%. ADGC genotype data was generated using 3 major genotyping platforms: Affymetrix 6.0, Illumina Human610, and Illumina Human660.

We first tried to see the feasibility of HP imputation using the directly genotyped data. We extracted the *HP* region genotype markers of the three genotyping platforms from the GTEx genotype data, respectively. We then performed HP imputation using the extracted data, respectively. We found the NA rate was high (Table S1).

Therefore, we then tried a TOPMed background imputation before the HP imputation (see manuscript Method for details). Using this method, we successfully reduced the NA rate (Table S2). In the *HP2* allele, exons 5 and 6 are repeated from exons 3 and 4. Ideally, the junction of exons 4&5 should be unique to *HP2*. By plotting the imputed *HP* genotypes and the exons 4&5 junction count, we see no clear error from this imputation approach (Figure S1) despite some potential alignment errors.

#### 3. Modeling Strategy and Exploring the APOEε2-3-4 Assumption

*Representing Main Effects of HP and APOE*

The complexity of interaction models can impact both the statistical power for detecting interactions and the interpretation of interaction effects. For bi-allelic variants such as the *HP* structural variant in this study, an additive or dose-per-allele model of genotypes has been previously shown to optimize power to detect an additive or dominant effect [11]. We also evaluated dominant and recessive models of *HP2* and found no significant effects. As such, we used an additive encoding of the *HP* alleles throughout all models.

There are three APOE-*ε* alleles (haplotypes of two linked SNVs) and thus six possible genotypes *ε2ε2*, *ε2ε3*, *ε2ε4*, *ε3ε3*, *ε3ε4*, *ε4ε4*. Multiple studies have confirmed that the *ε2* is protective relative to *ε3* and *ε4* increases risk relative to *ε3*. APOE-*ε* genotypes are often split into separate components, modeling an additive effect of either the *ε2* or *ε4* allele relative to *ε3ε3* individuals to provide a dose-per-allele interpretation of the genotypes (X decrease in AD risk per *ε2* allele, or X increase in AD risk per *ε4* allele). Other possibilities for *APOE* allele encodings are a categorical definition (fitting a different effect for each possible genotype which requires five coefficients).

Another possibility for representing the *APOE* genotypes is to encode a dose-by-allele effect which assumes a uniform increase in risk from *ε2* to *ε3*, and from *ε3* to *ε4*. This encoding is equivalent to an additive encoding, but is established for each of the two *APOE* alleles rather than for the genotype combinations of a biallelic SNP (as with *HP* described above). Because this encoding is on an allelic level, two coefficients are required (one for each allele), and an interaction term between the two alleles is required to account for genotype effects. This encoding requires three coefficients.

*Representing Interactions between HP and APOE*

Interactions among genotypes within a regression model framework can fully decompose the effects by encoding each genotype combination as a separate category, which for a bi-allelic by tri-allelic variant would result in 18 possible genotype combinations and would require fitting 17 coefficients.

To reduce the degrees of freedom in the model, the genotype effects can be encoded in a way that constrains the model as outlined in the main effects section above. For example, we could assume an additive effect of the *HP* genotype – this would require fitting one coefficient for the main dose-response effect for each *HP* non-reference allele (0, 1, or 2) in the context of a selected reference APOE genotype (i.e. *ε3ε3*). The main effects of this model would thus require six coefficients (one for HP and one for each non-reference category of *APOE* genotype). We would then include five additional interaction terms (one for each non-reference *APOE* genotype) that would estimate the change in dose-response per non-reference *HP* allele (relative to the *ε3ε3*) that is observed within each *APOE* genotype. The resulting model would contain eleven coefficients.

To further reduce the degrees of freedom expended, we could assume a dose-by-allele encoding of the *APOE* genotypes along with an additive encoding of the *HP* genotypes. Representing these main effects would require four coefficients (one for *HP*, two for each *APOE* allele, and one interaction for the *APOE* genotype effect). Three terms are required to represent the interaction of *HP* and *APOE*, one for the interaction of *HP* with each *APOE* allele, and one three-way interaction for the effect of *HP* by the *APOE* genotype effect. The main reason of choosing the *APOEε2-3-4* effect instead of running analysis using *APOE* alleles as factors with *ε3* as the reference allele is to reduce model complexity and gain model interpretability.

*Sensitivity Tests for the APOEε2-3-4 Assumption*
We examined a model where *APOE* genotypes and *HP* genotypes and their interactions were considered as different categories of a factor variable (*APOEε33 - HP1/HP2* as reference) and the significant effects from the stratified models detected from that model were qualitatively similar (Table S3).

Because we know from prior studies that the effect of *ε2*-to-*ε3* is not equivalent to *ε3*-to-*ε4*, it is possible that our assumption of a linear effect biases our interaction analysis [12], [13]. To explore the impact of this linear assumption, we performed a sensitivity analysis by adding another coefficient for the *ε3* status (*ε3*: 1, *ε2* and *ε4*: 0) in our primary model (Table S4). We observed a significant negative effect estimate of *ε3* from both *APOEε2* and *APOEε4* background models, suggesting that the *ε3* effect is closer to *ε2* than *ε4* (Table S5). Meanwhile, the interaction effect of *HP* and *APOEε2-3-4* remained significant in all models. We also examined dominant encodings of the APOE alleles – both *ε2* + *ε3* vs *ε4* and *ε2* vs *ε3* + *ε4.* We found that the *APOEε2-3-4* additive encoding has the best Akaike information criterion (AIC) over both of the dominant encodings – suggesting a better model performance, thus this is the model we have reported as our primary finding.

#### 4. Analysis of *HP* eQTL

The rs2000999 was shown to be associated with the RNA expression of HP that is independent of the *HP* SV [14]. In order to examine if the *HP* SV effect on AD risk is driven by expression difference, we repeated our main model with rs2000999 genotype included as a predictor. We found this eQTL has no impact on our association results (Table S6).

#### 5. African American (AA) Individuals' HP Imputation

The AA cohorts include the ACT, ADCs, CHAP, MIRAGE, NIA-LOAD, TARCC, the Indianapolis Ibadan Dementia Study (Indianapolis), Research in African-American Alzheimer's Disease Initiative (REAAADI) [15], GenerAAtions [16]. All subjects were recruited under protocols approved by the appropriate Institutional Review Boards.

Detailed information about the cohorts can be found at [16]. The detailed genotyping and phenotyping information can be found at [16]

The HP imputation of AA is very similar to the HP imputation European individuals described in the main text method. The only difference is that, since AA genome has an admix part of European ancestry [17], we imputed the *HP* genotypes in AA using both African and European references, as prior work has shown “cosmopolitan” reference panels increase imputation performance [18].

#### 6. Meta-Analysis of European and AA Data

After imputing the *HP* genotypes in the AA dataset, we first repeated the European cohort analysis. No significant effect of interest on AD risk was found in the logistic regression in AA data alone (Table S7), though the effect estimates are in consistent directions with EUR models (Table S8, S9). We conducted random effects meta-analysis of the logistic regression models in the main text using R *metafor* library [19], and find that the meta-analysis retains statistical significance.

#### Supplement Tables:

Table S1. Direct HP Imputation Genotype Count for GTEx Individuals Using Marker Sets of Affymetrix 6.0, Illumina Human610, and Illumina Human660, Respectively.

| | *HP1/HP1* | *HP1/HP2* | *HP2/HP2* | NA |
|---|---|---|---|---|
| Affymetrix 6.0 | 109 | 254 | 182 | 90 |
| Illumina Human610 | 103 | 256 | 180 | 96 |
| Illumina Human660 | 102 | 258 | 181 | 94 |

Table S2. Direct HP Imputation Genotype Count for GTEx Individuals Using Marker Sets of Affymetrix 6.0, Illumina Human610, and Illumina Human660 with TOPMed Background Imputation, Respectively.

| | *HP1/HP1* | *HP1/HP2* | *HP2/HP2* | NA |
|---|---|---|---|---|
| Affymetrix 6.0 | 127 | 295 | 189 | 24 |
| Illumina Human610 | 127 | 295 | 187 | 26 |
| Illumina Human660 | 127 | 296 | 187 | 25 |

Table S3. Logistic Regression Model on AD Risk Using Full Data (N=22,651) with All *HP-APOE* Interaction Groups. “Age” represents the age at onset for AD cases and age at last visit for controls.

| Variable | OR | 95%CI | p_value |
|---|---|---|---|
| (Intercept) | 2.42 | (1.80, 3.25) | 5.32e-09 |
| *HP1/HP1 + APOEε2ε2* | 0.34 | (0.10, 1.15) | 0.082 |
| *HP1/HP1 + APOEε2ε3* | 0.55 | (0.41, 0.74) | 7.81e-05 |
| *HP1/HP1 + APOEε2ε4* | 3.35 | (2.15, 5.22) | 8.46e-08 |

| | | | |
|---|---|---|---|
| *HP1/HP1 + APOEε3ε3* | 1 | (0.88, 1.12) | 0.944 |
| *HP1/HP1 + APOEε3ε4* | 3.46 | (3.02, 3.98) | 1.02e-69 |
| *HP1/HP1 + APOEε4ε4* | 8.88 | (6.42, 12.29) | 1.28e-39 |
| *HP1/HP2 + APOEε2ε2* | 0.42 | (0.17, 1.02) | 0.055 |
| *HP1/HP2 + APOEε2ε3* | 0.61 | (0.52, 0.73) | 1.83e-08 |
| *HP1/HP2 + APOEε2ε4* | 2.34 | (1.83, 2.98) | 7.38e-12 |
| *HP1/HP2 + APOEε3ε4* | 3.4 | (3.11, 3.73) | 2.39e-153 |
| *HP1/HP2 + APOEε4ε4* | 8.88 | (7.26, 10.87) | 1.11e-99 |
| *HP2/HP2 + APOEε2ε2* | 0.82 | (0.39, 1.72) | 0.599 |
| *HP2/HP2 + APOEε2ε3* | 0.61 | (0.51, 0.73) | 3.31e-08 |
| *HP2/HP2 + APOEε2ε4* | 2.07 | (1.56, 2.75) | 4.88e-07 |
| *HP2/HP2 + APOEε3ε3* | 0.98 | (0.90, 1.07) | 0.636 |
| *HP2/HP2 + APOEε3ε4* | 3.41 | (3.10, 3.76) | 2.57e-138 |
| *HP2/HP2 + APOEε4ε4* | 11.32 | (8.96, 14.30) | 6.09e-92 |
| Sex: Female | 0.98 | (0.93, 1.04) | 0.569 |
| Age | 0.98 | (0.98, 0.98) | 1.08e-25 |
| PC1 | 0.69 | (0.28, 1.70) | 0.423 |
| PC2 | 0.45 | (0.18, 1.10) | 0.08 |
| PC3 | 1.68 | (0.68, 4.20) | 0.264 |

Table S4. Logistic Regression Model on AD Risk with Full Data (N=22,651). *HP2* was included as the count of *HP2* alleles. "*APOE1*" and "*APOE2*" represents the *APOEε2-3-4* effects from the 2 copies of *APOE* alleles, respectively. "*APOE: if ε3*" represents an independent effect of levels 0 and 1, indicating if the *APOE* allele is *ε3* or not. "Age" represents the age at onset for AD cases and age at last visit for controls.

| Variable | OR | 95%CI | p_value |
|---|---|---|---|
| (Intercept) | 0.03 | (0.00, 0.26) | 0.001 |
| *HP2* | 7.26 | (1.75, 30.12) | 0.006 |
| *APOE1 ε2-3-4* | 4.67 | (1.82, 12.02) | 0.001 |
| *APOE2 ε2-3-4* | 5.19 | (2.48, 10.86) | 1.205e-05 |
| *APOE1*: if ε3 | 0.67 | (0.53, 0.85) | 8.774e-04 |
| *APOE2*: if ε3 | 0.69 | (0.42, 1.13) | 0.138 |
| Sex: Female | 0.98 | (0.93, 1.04) | 0.550 |
| Age | 0.98 | (0.98, 0.98) | 2.934e-25 |
| PC1 | 0.68 | (0.28, 1.67) | 0.399 |
| PC2 | 0.46 | (0.19, 1.13) | 0.090 |
| PC3 | 1.69 | (0.68, 4.21) | 0.263 |
| *HP2* x *APOE1 ε2-3-4* | 0.38 | (0.19, 0.74) | 0.004 |
| *HP2* x *APOE2 ε2-3-4* | 0.47 | (0.28, 0.78) | 0.003 |
| *APOE1 ε2-3-4* x *APOE2 ε2-3-4* | 0.70 | (0.50, 0.99) | 0.041 |

| | | | |
|---|---|---|---|
| *HP2* x *APOE1*: if ε3 | 1.01 | (0.86, 1.20) | 0.867 |
| *HP2* x *APOE2*: if ε3 | 0.81 | (0.58, 1.14) | 0.232 |
| *APOE2*: if ε3 x *APOE2*: if ε3 | 1.09 | (0.69, 1.72) | 0.721 |
| *HP2* x *APOE1 ε2-3-4* x *APOE2 ε2-3-4* | 1.45 | (1.14, 1.85) | 0.002 |
| *HP2* x *APOE1*: if ε3 x *APOE2*: if ε3 | 1.20 | (0.87, 1.65) | 0.278 |

Table S5. Logistic Regression Models on AD Risk Stratified by *APOE* Status. *HP2* was included as the count of *HP2* alleles. "*APOE: if ε3*" represents an independent effect of levels 0 and 1, indicating if the *APOE* allele is *ε3* or not. "Age" represents the age at onset for AD cases and age at last visit for controls.

| | ***APOEε2* (N=2,515)** | | | ***APOEε3* (N=20,261)** | | | ***APOEε4* (N=9,949)** | | |
|---|---|---|---|---|---|---|---|---|---|
| Variable | OR | 95%CI | p_value | OR | 95%CI | p_value | OR | 95%CI | p_value |
| (Intercept) | 0.24 | (0.06, 0.94) | 0.041 | 1.27 | (0.87, 1.85) | 0.212 | 1.27 | (0.87, 1.85) | 0.212 |
| *HP2* | 1.67 | (0.83, 3.37) | 0.151 | 1.03 | (0.89, 1.21) | 0.664 | 1.03 | (0.89, 1.21) | 0.664 |
| *APOEε2-3-4* | 3.36 | (1.93, 5.86) | 1.961e-05 | 2.45 | (2.16, 2.76) | 2.267e-46 | 2.45 | (2.16, 2.76) | 2.267e-46 |
| *APOE*: if ε3 | 0.62 | (0.34, 1.13) | 0.117 | 0.71 | (0.62, 0.83) | 8.320e-06 | 0.71 | (0.62, 0.83) | 8.320e-06 |
| Sex: Female | 1.01 | (0.84, 1.21) | 0.916 | 0.98 | (0.93, 1.04) | 0.585 | 0.98 | (0.93, 1.04) | 0.585 |
| Age | 0.99 | (0.98, 1.00) | 0.232 | 0.98 | (0.98, 0.99) | 1.256e-21 | 0.98 | (0.98, 0.99) | 1.256e-21 |
| PC1 | 3.42 | (0.23, 50.60) | 0.371 | 0.60 | (0.24, 1.51) | 0.280 | 0.60 | (0.24, 1.51) | 0.280 |
| PC2 | 0.05 | (0.00, 0.71) | 0.026 | 0.59 | (0.23, 1.50) | 0.270 | 0.59 | (0.23, 1.50) | 0.270 |
| PC3 | 6.67 | (0.37, 118.98) | 0.197 | 1.76 | (0.69, 4.53) | 0.238 | 1.76 | (0.69, 4.53) | 0.238 |
| *HP2* x *APOEε2-3-4* | 0.70 | (0.48, 1.01) | 0.056 | 0.98 | (0.90, 1.07) | 0.650 | 0.98 | (0.90, 1.07) | 0.650 |
| *HP2* x *APOE*: if ε3 | 0.88 | (0.59, 1.32) | 0.543 | 0.97 | (0.88, 1.08) | 0.614 | 0.97 | (0.88, 1.08) | 0.614 |

Table S6. Logistic Regression Models on AD Risk with Full Data. *HP2* was included as the count of *HP2* alleles. "*APOE1*" and "*APOE2*" represents the *APOEε2-3-4* effects from the 2 copies of *APOE* alleles, respectively. "rs2000999" represents the genotype of the HP eQTL rs2000999. "Age" represents the age at onset for AD cases and age at last visit for controls.

| Variable | OR | 95%CI | p_value |
|---|---|---|---|
| (Intercept) | 0.03 | (0.01, 0.13) | 5.17e-06 |

| | | | |
|---|---|---|---|
| *HP2* | 4.28 | (1.47, 12.42) | 0.008 |
| *APOE1 ε2-3-4* | 2.94 | (1.38, 6.26) | 0.005 |
| *APOE2 ε2-3-4* | 4.93 | (2.74, 8.87) | 1.06e-07 |
| Sex: Female | 0.97 | (0.92, 1.03) | 0.4 |
| Age | 0.98 | (0.98, 0.98) | 1.86e-25 |
| PC1 | 0.83 | (0.33, 2.07) | 0.693 |
| PC2 | 0.44 | (0.18, 1.10) | 0.08 |
| PC3 | 1.83 | (0.71, 4.68) | 0.209 |
| rs2000999: 0/1 | 0.97 | (0.90, 1.04) | 0.345 |
| rs2000999: 1/1 | 1.01 | (0.86, 1.18) | 0.94 |
| *HP2* x *APOE1 ε2-3-4* | 0.48 | (0.28, 0.83) | 0.008 |
| *HP2* x *APOE2 ε2-3-4* | 0.53 | (0.35, 0.81) | 0.003 |
| *APOE1 ε2-3-4* x *APOE2 ε2-3-4* | 0.83 | (0.62, 1.10) | 0.191 |
| *HP2* x *APOE1 ε2-3-4* x *APOE2 ε2-3-4* | 1.37 | (1.11, 1.68) | 0.003 |

Table S7. Effect of *HP* and both *APOE* alleles on AD risk in AA cohorts (N=4,617). Effect estimates are from a logistic regression model. *HP2* was included as the count of *HP2* alleles. “*APOE1*” and “*APOE2*” represents the *APOEε2-3-4* effects from the 2 copies of APOE alleles, respectively. “Age” represents the age at onset for AD cases and age at last visit for controls.

| Variable | OR | 95%CI | p_value |
|---|---|---|---|
| (Intercept) | 0.02 | (0.00, 0.27) | 0.003 |
| *HP2* | 2.42 | (0.32, 18.12) | 0.39 |
| *APOE1 ε2-3-4* | 1.01 | (0.29, 3.52) | 0.984 |
| *APOE2 ε2-3-4* | 1.62 | (0.65, 4.05) | 0.303 |
| Sex: Female | 0.83 | (0.72, 0.96) | 0.014 |
| Age | 1.01 | (1.00, 1.02) | 0.005 |
| PC1 | 0.14 | (0.03, 0.76) | 0.023 |
| PC2 | 3.34 | (0.49, 22.71) | 0.217 |
| PC3 | 0.5 | (0.09, 2.77) | 0.43 |
| *HP2* x *APOE1 ε2-3-4* | 0.57 | (0.19, 1.67) | 0.302 |
| *HP2* x *APOE2 ε2-3-4* | 0.69 | (0.31, 1.52) | 0.356 |
| *APOE1 ε2-3-4* x *APOE2 ε2-3-4* | 1.3 | (0.81, 2.08) | 0.285 |
| *HP2* x *APOE1 ε2-3-4* x *APOE2 ε2-3-4* | 1.23 | (0.82, 1.86) | 0.318 |

Table S8. Random Effects Meta-Analysis Model Fit with Both European-Descent Individuals' and AA Data and Effects of Both *APOE* Alleles. *HP2* was included as the count of *HP2* alleles. "*APOE1*" and "*APOE2*" represents the *APOEε2-3-4* effects from the 2 copies of *APOE* alleles, respectively. "Age" represents the age at onset for AD cases and age at last visit for controls.

| Variable | OR | 95%CI | p_value |
|---|---|---|---|
| *HP2* Allele Count | 3.68 | (2.76, 4.59) | 0.005 |
| *APOE1 ε2-3-4* | 2.44 | (1.35, 3.53) | 0.11 |
| *APOE2 ε2-3-4* | 3.99 | (2.79, 5.2) | 0.024 |
| Sex: Female | 0.96 | (0.78, 1.14) | 0.626 |
| Age | 0.99 | (0.95, 1.02) | 0.426 |
| *HP2* x *APOE1 ε2-3-4* | 0.5 | (0.04, 0.97) | 0.004 |
| *HP2* x *APOE2 ε2-3-4* | 0.58 | (0.22, 0.94) | 0.003 |
| *APOE1 ε2-3-4* x *APOE2 ε2-3-4* | 0.91 | (0.45, 1.36) | 0.669 |
| *HP2* x *APOE1 ε2-3-4* x *APOE2 ε2-3-4* | 1.32 | (1.14, 1.5) | 0.002 |

Table S9. Random Effect Meta-Analysis Models Stratified by *APOE* status. The effect estimates are from random-effect meta-analysis of European and AA analyses. "Age" represents the age at onset for AD cases and age at last visit for controls.

| | ***APOEε2*** | | | ***APOEε3*** | | | ***APOEε4*** | | |
|---|---|---|---|---|---|---|---|---|---|
| Variable | OR | 95% CI | p_value | OR | 95% CI | p_value | OR | 95% CI | p_value |
| *HP2* | 1.85 | (1.3, 2.4) | 0.029 | 0.99 | (0.83, 1.15) | 0.909 | 0.73 | (0.46, 1 | 0.024 |
| *APOEε2-3-4* | 4.42 | (3.95, 4.9) | <0.0001 | 2.78 | (2.43, 3.13) | <0.0001 | 1.8 | (1.54, 2.07) | <0.0001 |
| Sex: Female | 0.95 | (0.78, 1.13) | 0.604 | 0.96 | (0.79, 1.13) | 0.619 | 0.96 | (0.88, 1.04) | 0.304 |
| Age | 1 | (0.96, 1.05) | 0.897 | 0.99 | (0.95, 1.03) | 0.483 | 0.97 | (0.94, 1) | 0.083 |
| *HP2* x *APOEε2-3-4* | 0.75 | (0.51, 0.98) | 0.016 | 0.99 | (0.93, 1.06) | 0.848 | 1.15 | (1.02, 1.28) | 0.033 |

#### Supplement Figure

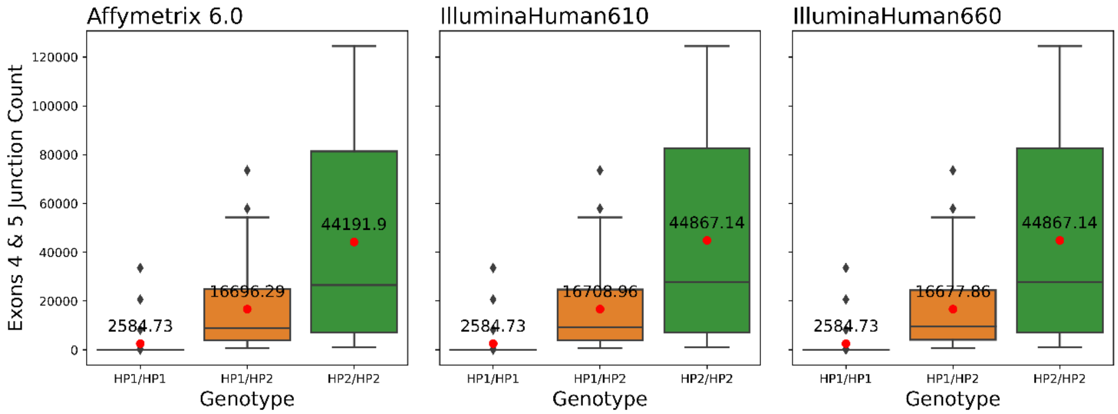


Figure S1. Boxplot of *HP* Exons 4 and 5 Junction Count vs. Imputed *HP* Genotypes for the GTEx Data Using Markers from Affymetrix 6.0, Illumina Human610, and Illumina Human660 Genotyping Platforms with TOPMed Background Imputation. Black bar in each box shows the median value of the group and the red dot shows the mean value.
